# PanCanSurvPlot: A Large-scale Pan-cancer Survival Analysis Web Application

**DOI:** 10.1101/2022.12.25.521884

**Authors:** Anqi Lin, Hong Yang, Ying Shi, Quan Cheng, Zaoqu Liu, Jian Zhang, Peng Luo

**Affiliations:** Department of Oncology, Zhujiang Hospital, Southern Medical University, Guangzhou, Guangdong, China; Department of Neurosurgery, Xiangya Hospital, Central South University, Changsha, Hunan, China; National Clinical Research Center for Geriatric Disorders, Xiangya Hospital, Central South University, Changsha, China; Department of Interventional Radiology, The First Affiliated Hospital of Zhengzhou University, Zhengzhou, Henan, China

**Keywords:** Survival Analysis, Pan-cancer, Prognosis, Tumor Prognostic Marker, Shiny, Web Application

## Abstract

The identification of reliable tumor prognostic markers can help clinicians and researchers predict tumor development and patient survival outcomes more accurately, which plays a vital role in clinical diagnosis, treatment effectiveness assessment, and prognostic evaluation. Existing web tools supporting online survival analysis are gradually failing to meet the increasing demands of researchers in terms of the dataset size, richness of survival analysis methods, and diversity of customization features. Therefore, there is an urgent need for a large-scale, one-stop pan-cancer survival analysis web server. We developed PanCanSurvPlot (https://smuonco.shinyapps.io/PanCanSurvPlot/), a Shiny web tool that has incorporated a total of 215 cancer-related datasets from the GEO and TCGA databases, covering nearly 100,000 genes (mRNAs, miRNAs, and lncRNAs), approximately 45,000 samples, 51 different cancer types, and 13 different survival outcomes. The website also provides two cutoff methods based on median and optimal cutpoints. All survival analysis results from the log-rank test and univariate Cox regression are presented in a clear and straightforward summary table. Finally, users can customize color schemes and cutpoint levels to quickly obtain high-quality Kaplan-Meier survival plots that meet publication requirements.

## Introduction

Tumor markers are specific measures that reflect a tumor’s molecular or histopathological characteristics. The detection of marker levels in patients prior to treatment is widely used to assess the occurrence of clinical events, disease recurrence or progression, and survival outcomes, thereby helping clinicians and researchers guide the monitoring of cancer patients and make decisions about personalized treatment[1–3]. Tumor prognostic markers are currently being studied in numerous areas, including genomics[4,5], transcriptomics[6–8], proteomics[9,10], metabolomics[11], and epigenomics[12,13]. Among these, the investigation of transcriptomic prognostic markers has been a hot area of research in the last decade owing to the rapid development of microarray and high-throughput sequencing technologies, which researchers have used to uncover a large number of potential molecular mechanisms of gene expression differences leading to disease development and clinical applications. For instance, based on evidence from numerous randomized controlled trials and clinical trials, patients with HER2-negative, ER-positive, peripheral lymph node-negative early invasive breast cancer who are younger than 50 years old and have expression-based 21-gene scores in the range of 16–25 have significantly longer invasive disease-free survival after chemotherapy[14,15]. The well-established Oncotype DX test is therefore used clinically to conveniently calculate 21-gene scores to assess patients’ suitability for adjuvant endocrine therapy and chemotherapy[14,15].

Several publicly available databases storing pan-cancer gene expression profile data, such as The Cancer Genome Atlas (TCGA)[16] and Gene Expression Omnibus (GEO)[17,18], provide us with detailed and reliable transcriptomic data accompanied by clinical and survival information. However, for clinicians and researchers, the direct utilization of transcriptomic data for research and clinical guidance is still a significant obstacle due to the need for more expertise related to array/sequencing and the ability to analyze data using programming languages[19]. Therefore, a web-based analysis tool that can accurately and efficiently assess tumor prognostic markers is greatly needed, which can lower the technical threshold and improve the efficiency of cancer research. Numerous web tools exploring prognostic markers in tumor transcriptomics have emerged, such as GENT2[20], GEPIA2[21], PROGgeneV2[22], OncoLnc[23], and Kaplan-Meier Plotter[24]. These web tools integrate gene expression profiling with clinical survival data and greatly facilitate users’ exploration of tumor prognostic markers through a simple and easy-to-understand interface and operation.

However, the above applications still have some unresolved or pending issues regarding the function and practical application of web tools. For example, in terms of the number of included datasets and samples, most of these web tools only contain partial datasets from the TCGA or GEO databases and do not instantly update the latest datasets and include more prosperous and comprehensive tumor types. In terms of the richness of survival data, often only limited types of survival information are provided, and there is still a large amount of valuable survival information that needs to be better utilized. More importantly, regarding survival analysis algorithms, some web tools only provide log-rank tests and visualization based on the median gene expression cutoff, which fails to meet the needs of researchers for diverse survival analysis methods and parameters. A web tool that incorporates extensive pan-cancer transcriptomic data and clinical survival information and provides multiple survival analysis methods with custom graphing capabilities would better facilitate the needs of oncology researchers and clinicians for analysis.

Therefore, we are developing a web tool, PanCanSurvPlot (https://smuonco.shinyapps.io/PanCanSurvPlot/), based on the existing R shiny framework, focusing on the visualization of pan-cancer survival analysis results for a broad group of biomedical researchers (with or without programming skills). We intend to collect tumor-related datasets with complete detailed clinical survival information and transcriptomic data, covering as many cancer types and types of survival information as possible, retrieved from several large publicly available databases, including GEO and TCGA. Standard survival analysis methods will be provided, including the log-rank test, univariate Cox analysis, and custom visualization features. In addition, the simple and easy-to-understand interface and the user-friendly search function will better serve users’ needs for cross-sectional comparisons of the prognostic value of the same gene in different cancers or datasets. Various customization functions, publication-ready visualizations, and detailed raw data information will help users conduct further in-depth analysis.

## Materials and Methods

### Data Collection and Preprocessing

PanCanSurvPlot collected tumor-related microarray or high-throughput sequencing datasets from the GEO and TCGA databases. The inclusion criteria for datasets were as follows: (i) expression profile dataset (microarray or high-throughput sequencing) of human tumor samples; (ii) available corresponding detailed clinical and survival information (including survival time and survival status); and (iii) number of tumor samples of at least 30. The exclusion criteria for datasets were as follows: (i) the sample source was not human; (ii) the sample source was nontumor tissue; (iii) the expression profile data were abnormal or did not meet the analysis requirements; (iv) clinical information and survival information were abnormal or missing; and (v) the number of tumor samples was too small (<30). For datasets containing multiple platforms, we treated the data from each platform as independent datasets. Finally, the corresponding expression profile and clinical data of 182 GEO datasets were downloaded from the GEO website and GEOquery package[25]. The corresponding expression profile and clinical data of 33 TCGA datasets were downloaded from the UCSC XENA platform. All downloaded expression profiles were double-checked for normalization and log2 transformation. All high-throughput datasets were prenormalized, while microarray datasets without normalization were standardized using the limma package[26]. As the unit for survival data, we standardized the survival time to months. Samples missing corresponding survival information were removed before proceeding to the next step of the analysis. An overview of the included datasets is provided in Supplementary Table 1.

### Survival Analysis and Visualization

For each dataset, gene expression was divided into high and low groups based on the median (or optimal cutoff value). As described in the previous literature[27], group sample sizes that are too small would result in unreliable analysis results. Therefore, when using the median as the cutoff value, we excluded genes with extremely imbalanced grouping (sample size below 5% of the total and above 95% of the total under either grouping). Implementing the optimal cutoff value calculation relies on the surv_cutpoint function in the survminer package (v0.4.9), with the default setting that requires any subgroup’s sample size to be no less than 5% of the total. For survival analysis, survival curves were plotted using the Kaplan-Meier method, and the log-rank test and univariate Cox proportional hazards regression model were applied to assess the effect of individual gene expression on the clinical outcome, implemented by the survfit and coxph functions within the surv package (v3.4-0). Hazard ratios (HRs), 95% confidence intervals (95% CIs), and p values under different grouping criteria and survival analysis methods were calculated accordingly. Based on the results, we integrated a table of the survival analysis results for each gene in the corresponding datasets into PanCanSurvPlot. The visualization of Kaplan-Meier survival analysis results was implemented through the survminer package.

### Web Server Implementation

PanCanSurvPlot (https://smuonco.shinyapps.io/PanCanSurvPlot/) was built mainly on R, a free and open-source programming tool for data analysis, statistics, and graphing; the R Shiny package for building interactive web tools; and shinyapps.io for architecting the server. HTML5, CSS, JavaScript, and other programming languages were also used to assist with web page construction and beautification. Figure 1 summarizes the workflow of PanCanSurvPlot.

**Figure 1:**
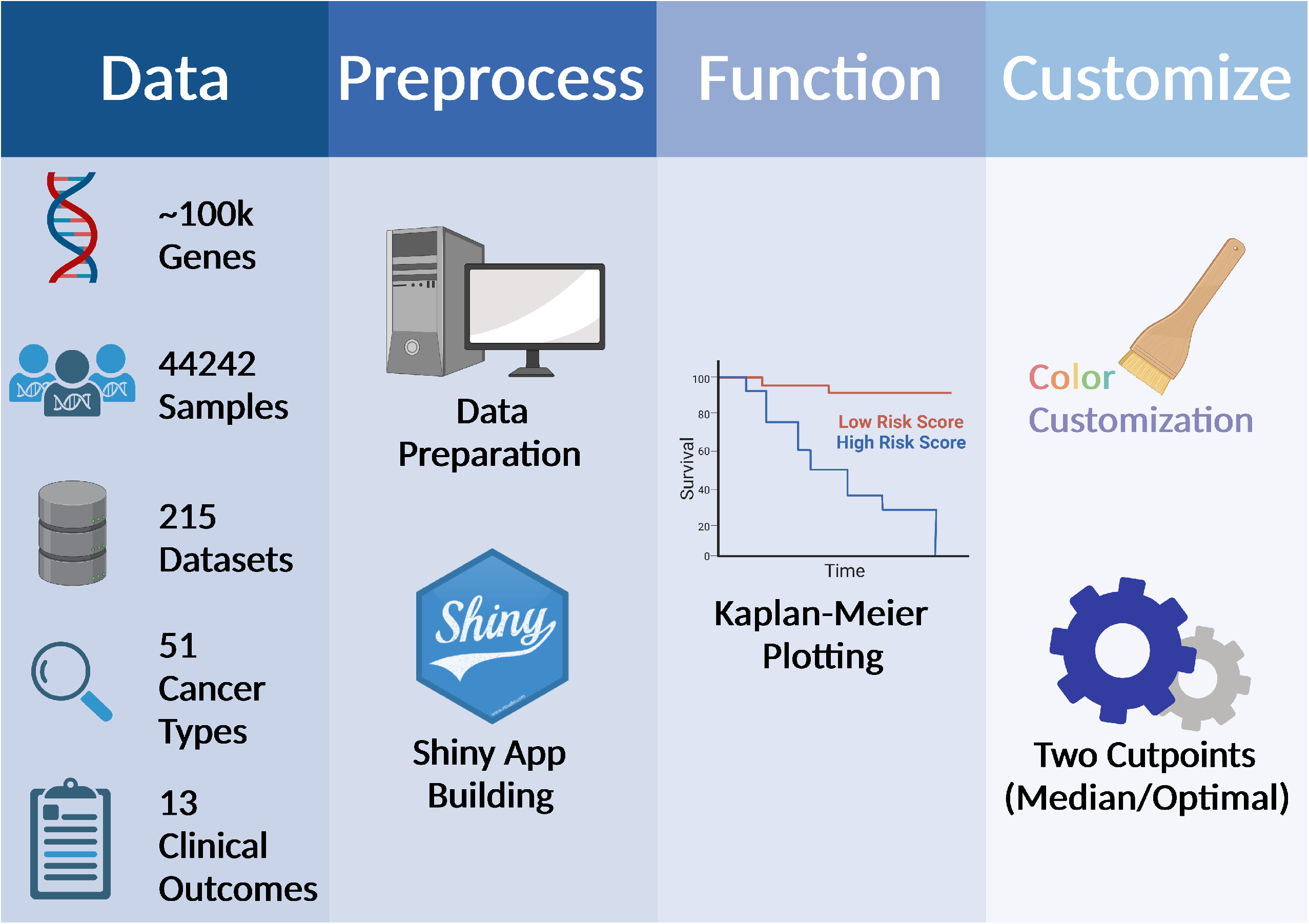
Workflow of PanCanSurvPlot PanCanSurvPlot (v1.1.0) currently incorporates 215 datasets from the TCGA and GEO databases. After data collection and preprocessing were completed, the survival analysis web tool PanCanSurvPlot was built through Shiny to provide high-quality Kaplan-Meier analysis, plotting, and rich customization features.

## Results

### Overview

PanCanSurvPlot is free and registration-free for all users. To date, PanCanSurvPlot (version 1.1.0) has incorporated 7,676,978 records distributed in 182 GEO datasets and 33 TCGA datasets, which cover approximately 100,000 genes (mRNAs, miRNAs, and lncRNAs), nearly 45,000 samples, 51 different cancer types, and 13 different survival outcomes. The main body of PanCanSurvPlot is divided into four sections: 1 HOME: Through simple and easy-to-understand text and pictures, we show users the function introduction, step-by-step guidance, and notification of the version update status of PanCanSurvPlot, so that users can quickly understand the design, purpose, and usage of the webpage. 2 PLOT: For the selected genes of interest, detailed survival analysis results and high-quality plots that meet publication requirements are easily accessible. 3 DATA: The interactive form helps users more easily access the overview of datasets included in PanCanSurvPlot and learn how to download them. 4 ABOUT: For any active queries, requests, or suggestions about PanCanSurvPlot, the contact details of the technical staff and FAQ can serve the users well. Figure 2 shows the interactive interface of the main panel of PanCanSurvPlot.

**Figure 2:**
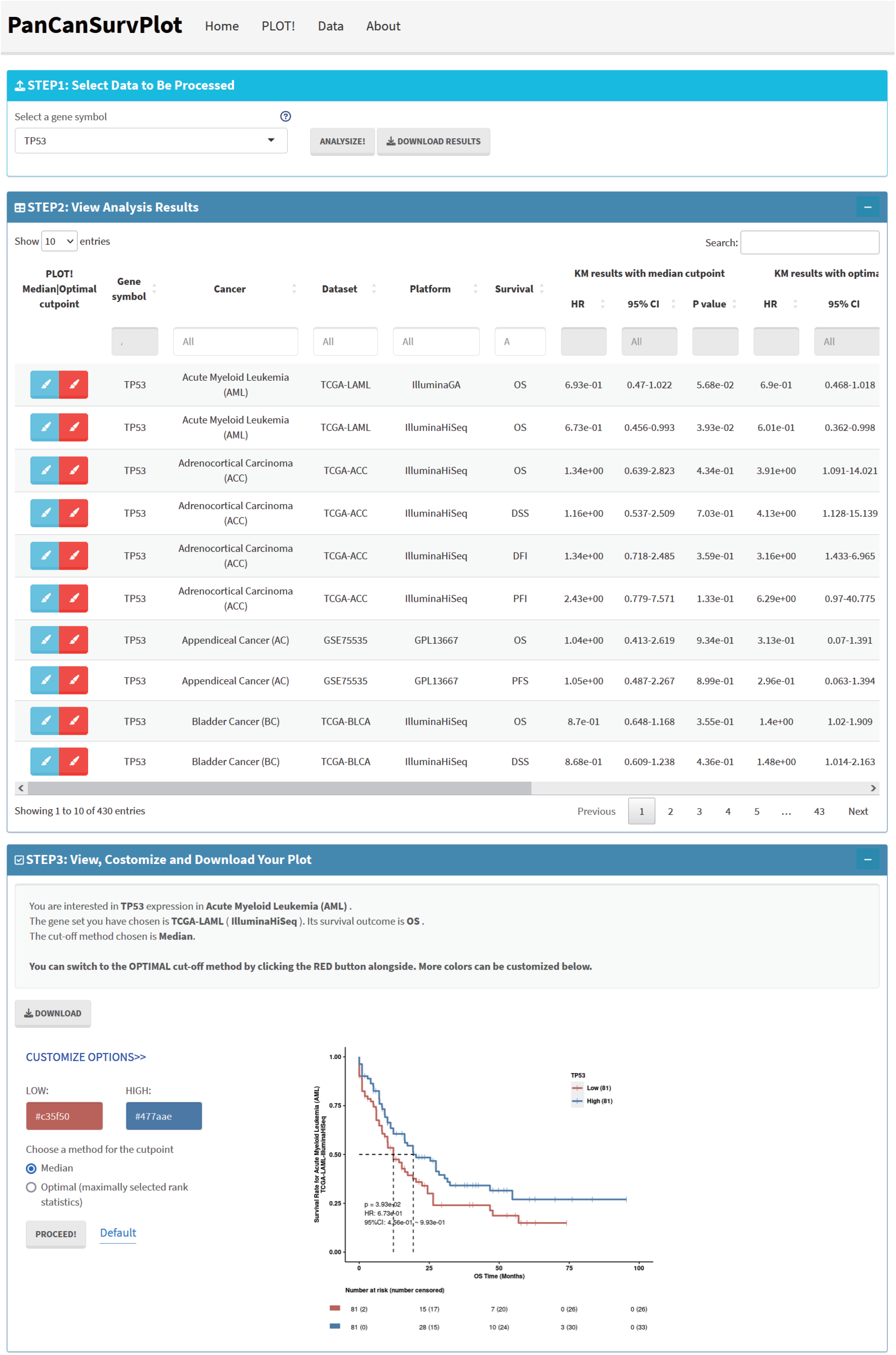
The main interface of PanCanSurvPlot The main interface of PanCanSurvPlot is divided into three parts: target gene selection, survival analysis result overview, and plotting and customization. PanCanSurvPlot provides a simple and straightforward survival analysis result overview table, high-quality plotting, and rich customization functions. The detailed explanatory text and the easy-to-understand interface are also user-friendly features of PanCanSurvPlot.

### Survival Analysis

PanCanSurvPlot enables survival analysis of single gene expression in the pan-cancer field. Under the PLOT page, users can select a specific gene of interest in the drop-down box or directly type it in. Nearly 100,000 genes, including mRNAs, miRNAs, and lncRNAs, are available for users to select. After clicking the submit button, users can quickly obtain the summary table of survival analysis results for that target gene in all datasets. In addition to basic information such as cancer type, dataset name, dataset platform, and type of survival outcomes, detailed statistical results (HR, 95% CI, and P value) from log-rank tests and univariate Cox regression are also provided in full. PanCanSurvPlot provides both median and optimal cutoff values in two grouping modes to meet the diverse needs of users for survival analysis. The table of survival analysis results (in CSV format) for the gene of interest can be quickly accessed via the download button.

The Kaplan-Meier plot, completed according to the user’s selection, is quickly displayed after clicking the corresponding plot button. The customization function on the side allows the user to make custom changes to the image’s color scheme and grouping method. Similarly, downloading high-resolution images in PDF format is also allowed. Taking the survival analysis of CD48 in the TCGA-BRCA dataset in the TCGA database and the GSE26304 breast cancer dataset in the GEO database as examples, the user obtains the results shown in Figures 3A and B. The CD48 high expression group in both datasets had a longer overall survival (OS), which validated the previous findings[28].

**Figure 3:**
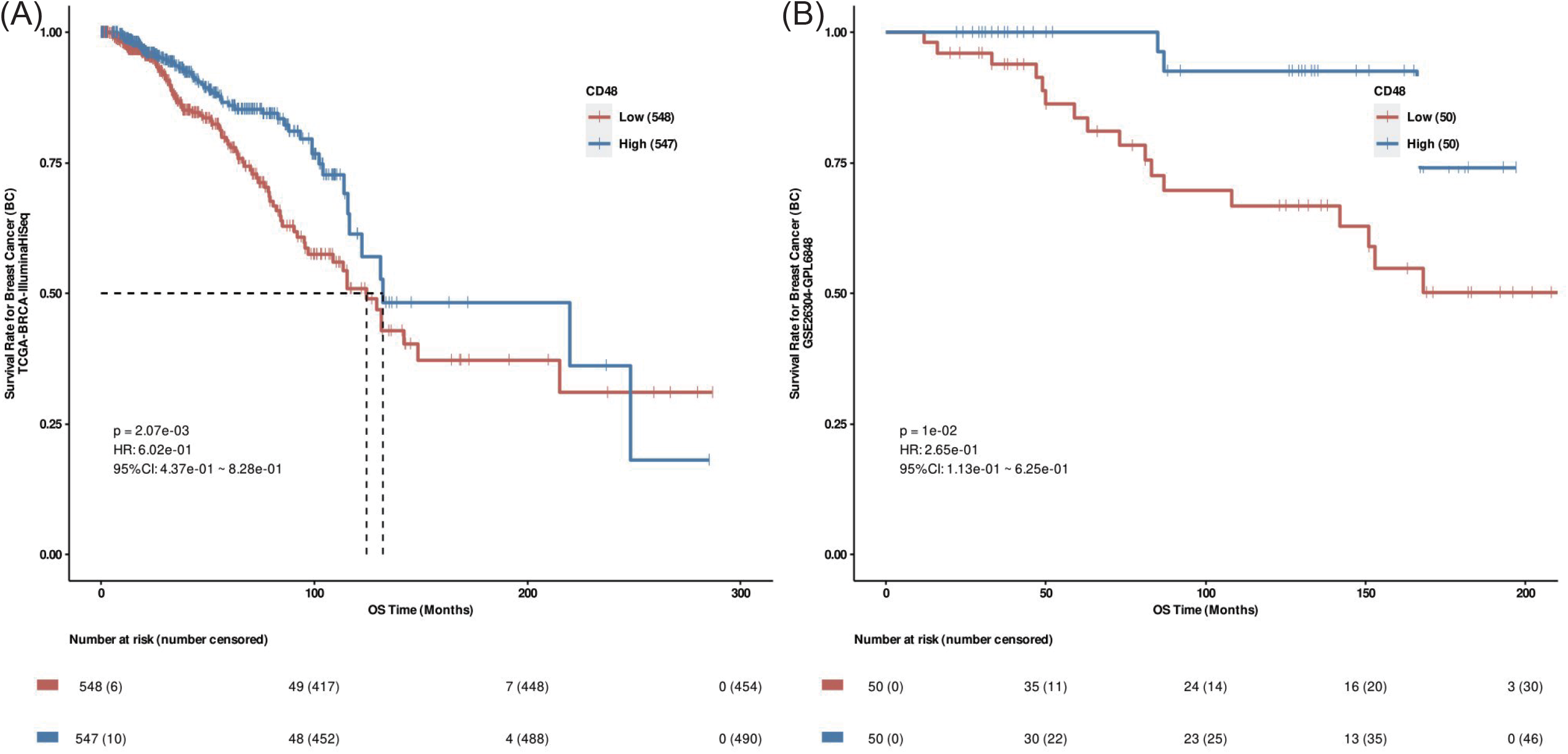
Survival analysis plotting results of PanCanSurvPlot Figure 3A and 3B show Kaplan-Meier plots of CD48 in TCGA-BRCA and GSE26304, respectively. P values were calculated by the log-rank test, with grouping based on the median CD48 expression. HR: hazard ratio, 95% CI: 95% confidence interval.

The DATA section provides detailed information on the included datasets for users to query quickly, including cancer types, datasets, platforms, survival outcomes, and links to the corresponding datasets. In addition, users can select numerous user-friendly features to meet the need to save data locally for further study, including copying to the clipboard, saving locally in CSV, EXCEL, and PDF formats, or printing.

### Documentation

The About section provides the contact details of the authors and technicians, and any feedback or feature requests for PanCanSurvPlot will be taken seriously. The records of all possible improvements to PanCanSurvPlot will be written in the update section of the home page afterward. In the FAQ section, we provide detailed answers to the questions that users may have. For example, the list of all genes covered by PanCanSurvPlot and the detailed description of the 13 survival outcomes are available here, which will better assist users in their oncology studies with PanCanSurvPlot.

## Discussion

PanCanSurvPlot is a large-scale interactive web tool dedicated to pan-cancer transcriptome survival analysis and visualization. The extensive tumor expression profile datasets and clinical survival information from the GEO and TCGA databases are fully utilized, making it easy for clinicians or researchers without sophisticated programming skills to manipulate to obtain the data needed for research. Through PanCanSurvPlot, users can quickly explore the impact of target genes on survival outcomes in different tumors, assisting clinicians and researchers in further investigating the mechanism of tumor development and improving clinical decision-making.

Compared with the emerging web-based survival analysis counterparts in recent years (Table 1), PanCanSurvPlot has different advantages in terms of the breadth of genes studied, the richness of cancer types, the size of datasets, the diversity of survival information, the comprehensiveness of analysis methods, and the user-friendliness of customization functions. In recent years, a large number of studies have suggested that lncRNAs are involved in regulating tumor development, and studying the expression characteristics of lncRNAs is one of the fundamental ways to develop further in-depth mechanistic studies[29,30]. Therefore, PanCanSurvPlot has its own unique value as a web-based tool covering mRNAs, miRNAs, and lncRNAs in cancer survival analysis. Compared with previous survival analysis tools that only utilize the TCGA and GEO databases and focus only on specific cancers, the inclusion of rich pan-cancer datasets and dozens of survival outcomes is undoubtedly another significant advantage of PanCanSurvPlot. More importantly, in the core module of survival analysis, PanCanSurvPlot tables visualize the prognostic value of target genes in multiple cancers in a clear way. The ability to simultaneously display the results of log-rank tests and univariate Cox analyses for target genes in different cancers based on two cutoff methods is a groundbreaking design that is not available in existing web tools, allowing users to obtain more intuitive and comprehensive statistical results for follow-up studies. In addition, the plotting customization feature will serve more users’ individual needs, allowing for quick access to survival analysis graphs that meet publication requirements.

**Table 1:** Comparison of PanCanSurvPlot with survival analysis web tool counterparts

|  | PanCanSurvPlot | GENT2 | GEPIA2 | PROGgeneV2 | OncoLnc | KM Plotter |
| --- | --- | --- | --- | --- | --- | --- |
| Data Type | mRNAs,<br>miRNAs,<br>LncRNAs | mRNAs | mRNAs,<br>miRNAs,<br>LncRNAs | mRNAs | mRNAs,<br>miRNAs,<br>LncRNAs | mRNAs,<br>miRNAs |
| Dataset* | 215 | Aggregated | 33 | 124 | 21 | Aggregated |
| Cancer Type* | 51 | 19 | 33 | 21 | 21 | 21 |
| Sample* | 44,242 | 3,038 | 9,664 | 28,503 | 7,982 | ~30,000 |
| Survival Outcome* | 13 | 4 | 2 | 5 | 1 | 4 |
| Optimal Cutpoint | YES | NO | NO | NO | NO | YES |
| Cox Analysis | YES | NO | NO | YES | YES | YES |
| Color Customization | YES | NO | YES | NO | NO | NO |
| Subgroup Analysis | NO | YES | YES | YES | NO | YES |
| Reference | - | [20] | [21] | [22] | [23] | [24] |

\* The number of datasets, cancer types, samples, and survival outcomes available in the survival analysis Aggregated: Datasets have been aggregated when conducting the survival analysis.

Admittedly, PanCanSurvPlot still has some limitations, which we intend to improve in the subsequent version updates. First, for the inclusion of datasets, PanCanSurvPlot currently only involves microarray and high-throughput sequencing datasets from the TCGA and GEO databases. There are still many potential target data in the additional materials of published articles, databases such as The Surveillance, Epidemiology, and End Results (SEER), Sequence Read Archive (SRA), ArrayExpress, and the continuously updated GEO database that we need to include. In addition, for survival analysis, PanCanSurvPlot currently cannot exclude the effect of primary clinical factors on survival, and the cutoff value can only be set to the median or the optimal cutoff value, which does not meet the possible needs of users to adjust the interference of clinical factors and customize the cutoff value of the survival analysis group. In this regard, our team will update PanCanSurvPlot in future versions to include more comprehensive cancer-related datasets and provide as many options as possible for survival analysis parameters. In the future, our team plans to conduct research on survival analysis web tools in fields other than transcriptomics, including genomics, proteomics, and epigenomics, to provide biomedical researchers with more comprehensive and extensive survival analysis assistance.

Overall, PanCanSurvPlot is a convenient, accessible, and intuitive web-based pan-cancer survival analysis web tool that can provide users, especially biomedical researchers who do not have programming skills, with the ability to explore the survival prediction results of transcriptomic markers for different tumors and different survival outcomes. By further extending the functionality of PanCanSurvPlot and continuously improving it based on user feedback, PanCanSurvPlot has the potential to become a solid part of routine analysis in the biomedical field.

## Supporting information

Supplementary Table 1: Overview of PanCanSurvPlot's incorporated dataset

---

Supplementary Table 1: Overview of PanCanSurvPlot’s incorporated dataset

## Acknowledgments

Not applicable.

## Author Contributions

All authors contributed to the study’s conception and design. AL, HY, and YS collected data, performed data analyses, and wrote the manuscript. QC, ZL, PL, and JZ reviewed the manuscript and supervised this work. All authors read and approved the final manuscript.

## Data Availability

All datasets presented here are generated by the TCGA Research Network (https://www.cancer.gov/tcga) and NCBI’s Gene Expression Omnibus (https://www.ncbi.nlm.nih.gov/geo). All other data generated and analyzed in this study are available from the corresponding authors upon reasonable request.

## Ethics Approval

Not applicable.

## Consent to Participate

Not applicable.

