## Supplementary Table 1: Overview of PanCanSurvPlot's incorporated dataset for "PanCanSurvPlot: A Large-scale Pan-cancer Survival Analysis Web Application"

| Cancer Type | Sample Counts | Datasets Count |
| --- | --- | --- |
| Acute Myeloid Leukemia | 173 | 1 |
| Adrenocortical Carcinoma | 79 | 1 |
| Appendiceal Cancer | 39 | 1 |
| Bladder Cancer | 897 | 7 |
| Breast Cancer | 11940 | 64 |
| Burkitt Lymphoma | 159 | 1 |
| Burkitt Lymphoma and Diffuse Large B Cell Lymphoma | 297 | 2 |
| Cervical Cancer | 602 | 2 |
| Cholangiocarcinoma | 36 | 1 |
| Chromophobe Renal Cell Carcinoma | 66 | 1 |
| Clear Cell Renal Cell Carcinoma | 944 | 1 |
| Colon Cancer | 1832 | 10 |
| Colorectal Cancer | 1506 | 15 |
| Cutaneous Melanoma | 468 | 1 |
| Diffuse Large B cell Lymphoma | 629 | 6 |
| Endometrial Carcinoma | 583 | 1 |
| Esophageal Carcinoma | 204 | 1 |
| Esophageal Squamous Cell Carcinoma | 179 | 2 |
| Follicular Lymphoma | 106 | 1 |
| Gastric Cancer | 1019 | 3 |
| Glioblastoma | 842 | 2 |
| Head and Neck Cancer | 641 | 3 |
| Hepatocellular Carcinoma | 727 | 5 |
| Invasive Lobular Carcinoma | 117 | 1 |
| Laryngeal Cancer | 165 | 2 |
| Liposarcoma | 140 | 1 |
| Lower Grade Glioma | 515 | 1 |
| Lung Adenocarcinoma | 756 | 2 |
| Lung Cancer | 717 | 3 |
| Lung Squamous Cell Carcinoma | 1182 | 7 |
| Melanoma | 210 | 1 |
| Mesothelioma | 87 | 1 |
| Neuroblastoma | 276 | 1 |
| Non Small Cell Lung Cancer | 1374 | 13 |
| Ovarian Cancer | 10463 | 22 |
| Pancreatic Cancer | 207 | 3 |
| Pancreatic Ductal Adenocarcinoma | 345 | 3 |
| Papillary Renal Cell Carcinoma | 290 | 1 |
| Papillary Thyroid Carcinoma | 505 | 1 |
| Paraganglioma and Pheochromocytoma | 179 | 1 |
| Prostate Adenocarcinoma | 497 | 1 |
| Prostate Cancer | 814 | 6 |
| Rectal Cancer | 413 | 3 |
| Renal Cell Carcinoma | 94 | 2 |
| Sarcoma | 259 | 1 |
| Testicular Germ Cell Cancer | 134 | 1 |
| Thymoma | 120 | 1 |
| Urothelial Cancer | 224 | 1 |
| Uterine Carcinosarcoma | 57 | 1 |
| Uterine Serous Carcinoma | 54 | 1 |
| Uveal Melanoma | 80 | 1 |
